# Biochemical analysis challenging Western blot analysis as validation step for antibodies intended for ELISA and Immunohistochemistry use

**DOI:** 10.1101/2022.02.14.480459

**Authors:** Yunyun Zhang, Wenfeng Zhang, Maozhou Yang, Jiandi Zhang

**Affiliations:** Yantai Quanticision Diagnostics, Inc., a division of Quanticision Diagnostics, Inc. of US Yantai, Shandong 264670, P. R. China

**Keywords:** FFPE, native, denatured, QDB, antigen, epitope, antigen-antibody interaction, ELISA, immunohistochemistry, antibody validation, Western blot

## Abstract

The relative contributions of the conformation and primary structure of an epitope to overall antigen-antibody interaction (AAI) at denatured, native or formalin fixed (FF) state were compared using six randomly chosen commercial antibodies using Quantitative Dot Blot (QDB) method. AAIs at native and FF states were found ranged 1.3 ∼ 10.2 and 0.5 ∼ 45.4 folds, respectively, over those at denatured state in cellular and tissue lysates. Using two antibodies against different epitopes of PYGL protein, we showed that PYGL levels in several types of tissues and cell lines were highly correlated (r=0.99 from Pearson, p<0.0001, n=25) when measured with these two antibodies at native state. Yet, one antibody was found to be nonspecific with one type of these tissues using Western blot analysis. These observations suggested that the conformation of an epitope may serve as dominant contributor of overall AAI at native state in general, regardless of linear or conformational epitopes. In many cases, it would override nonspecific interactions formed at denatured state to challenge Western blot analysis as a validation tool for antibodies intended for immunohistochemistry (IHC) and ELISA.

## Introduction

Both the conformational structure (shape) and the primary structure (sequence) of an epitope in an antigen are essential components of antigen-antibody interaction (AAI). An epitope is generally categorized as conformational or linear epitope in literature (1). The term “linear epitope” refers to those epitopes relying primarily on sequence, rather than shape, for AAI. Conversely, Conformational epitope refers to those epitopes recognized by the antibody primarily through shape rather than sequence. Not surprisingly, conformational epitope is also called discontinuous epitope while linear epitope is called continuous epitope.

A common assumption behind these definitions is that for continuous epitopes, the contributions from their shapes are minimal to AAI. Yet, there is limited attempt to validate this assumption. In other word, does the shape of a continuous epitope make any contribution to the overall AAI?

AAI is the pillar of immunodetection techniques including Enzyme linked Immunosorbent Assay (ELISA), Western blot analysis (WB), Reverse phase protein array and Immunohistochemistry (IHC). For WB, the antigen targets are usually fully denatured with all the conformational structure of the epitopes destroyed by the combined usage of heat, Sodium Dodecyl Sulfate (SDS), and reducing reagent [Dithiothreitol (DTT) or β-mercaptoethanol]. The term “linear epitope” fits its definition well in this case, as the sequence is the sole determinant of AAI at the denatured state.

On the other hand, the shape of the epitope is well preserved at native states. For antigens studied in IHC, they are extensively crosslinked by a fixative agent, most often by Formalin. While they are assumed at denatured state by many experts(2), there is little evidence that the conformation or shape of the epitope is affected by Formalin fixation (FF).

Recently, a high throughput immunoassay, Quantitative Dot Blot (QDB), was introduced to measure protein levels absolutely and quantitatively(3–7). This method is able to measure overall AAI at any states of the antigen including fully denatured, native and formalin fixed (FF) states. It thus can allow us to compare the performance of an antibody at all these three states objectively and quantitatively. Here we screened all the available antibodies in the laboratory with the sole requirement that these antibodies must be specific in WB analysis. We were able to identify six antibodies against FGFR_2_, CDK2, CDK4, PYGL, ATP5A and CAMKII proteins respectively. Their source information RRIDs were provided in Materials and Methods section(2). Among these six antibodies, four (CDK4, PYGL, ATP5A and CAMKII) are suggested to use for both Western blot analysis and immunohistochemistry (IHC) while the other two (FGFR_2_ and CDK2) are suggested for Western blot analysis only.

## Results and discussion

We defined lysates at the fully denatured form when they were heated in neutral lysis buffer at 85°C for 5mins in the presence of 50 mM DTT and 1%SDS. Those protein lysates in neutral lysis buffer without any treatment were defined as at native state. The FF state was defined as total protein lysates extracted from tissues pre-fixed in formalin solution for 16 hours without any other treatment. The mouse liver lysate was prepared accordingly, and was diluted from 0 to 1 μg/unit for QDB analysis. As shown in Fig. 1, all six antibodies recognized only one band of expected size in WB, confirming their specificity at denatured state. Unexpectedly, when measured the total AAIs with QDB method, we found all AAIs at native state were significantly higher than those at denatured state, ranging from 2.0 ∼ 8.8 over those at denatured state. Likewise, AAIs at FF state were also significantly higher than those at denatured states, ranging from 1.2 ∼ 45.4 folds over those at denatured state.

**Fig. 1.**
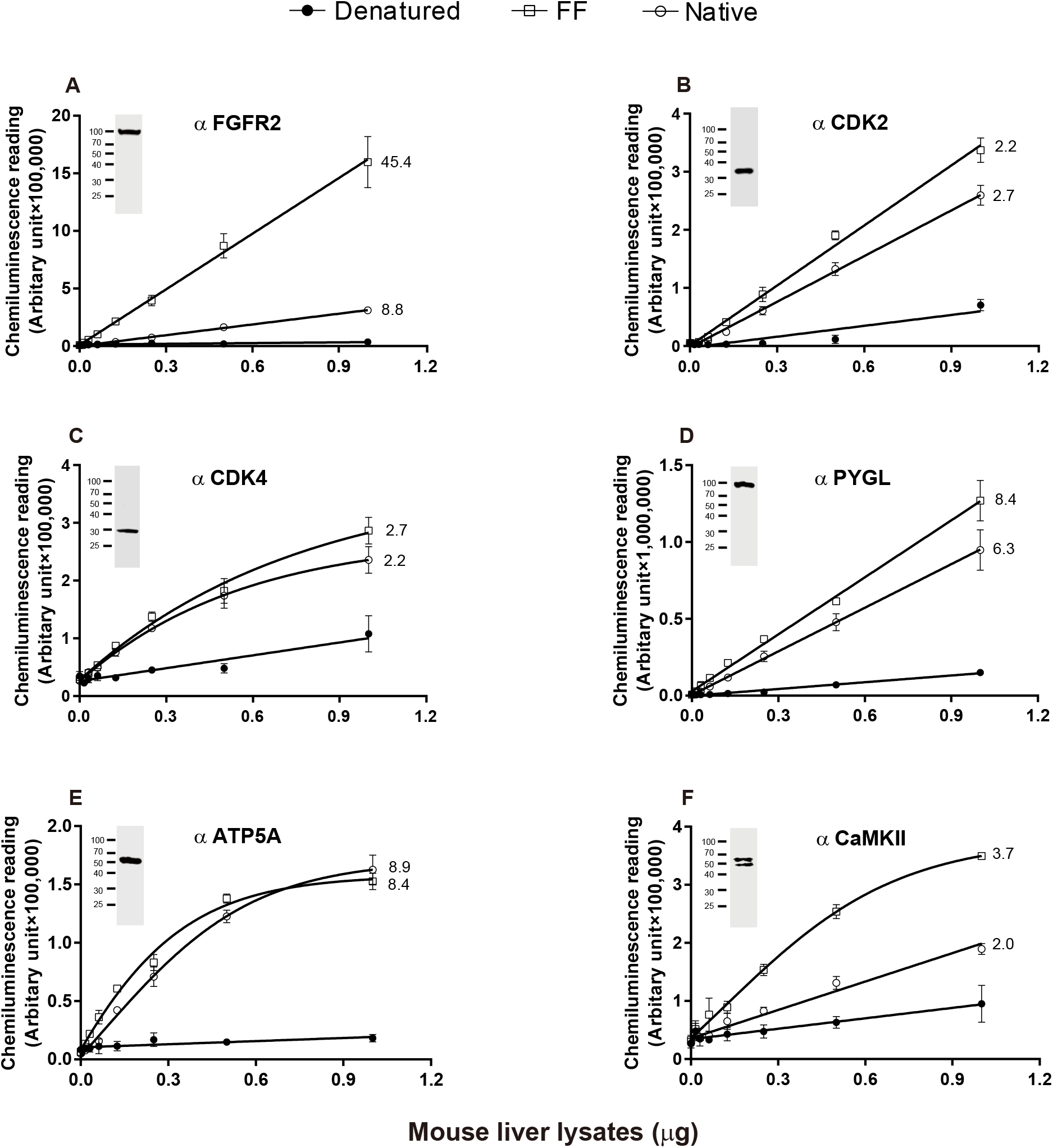
Comparison of performance of antibodies interacting with antigens at native, Formalin Fixed (FF) and denatured states in mouse liver. Whole tissue lysates from 6 mouse livers were extracted using lysis buffer containing Triton-X 100 buffer, and used directly as native state, or treated with 1% SDS, and 50 mM DTT for 5 mins at 85°C as denatured state. In a parallel experiment, the mouse liver tissues were fixed in 10% Formalin Solution for 16 hours and tissue lysates prepared as FF state. The total Antigen-antibody interactions (AAIs) at native, denatured and FF states for each antibody were plotted as in Fig. 1A to 1F respectively. The results from Western blot analysis of each antibody using denatured mouse liver lysates were inserted in Fig. 1A to 1F correspondingly.

We also prepared cellular lysates using HEK293 cells at all three states as described above, and diluted these lysates from 0 to 2 μg/unit. Again, all the antibodies recognized only one band of expected size in WB. We also observed that AAIs at native state were significantly higher, ranging from 1.3 ∼ 10.2, than those at denatured state. On the other hand, only three of the antibodies (FGFR2, ATP5A and CAMKII) recognized FF antigens significantly better than those of denatured ones. The other three antibodies recognized FF antigens either at similar level (CDK2 and PYGL) or worse (CDK4) than those of denatured state.

Even more surprisingly, the same CDK4 antibody was able to recognize FF antigen significantly better than denatured antigen in mouse liver, but not in HEK293 cells. This observation suggested that AAI interactions of CDK4 in mouse liver and human cell line may be different. Currently, we do not have a good explanation of this discrepancy (Fig. 1c and Fig. 2c). One putative explanation was that there might exist different cellular signaling pathway between mouse liver and HEK293 cell lines, where a unique factor in HEK293 cells might be “stapled” with CDK4 to block the access of anti-CDK4 antibody at FF state. Nonetheless, the difference between AAIs of native and FF states demonstrated that the FF state was indeed different from both native and denatured states.

**Fig. 2.**
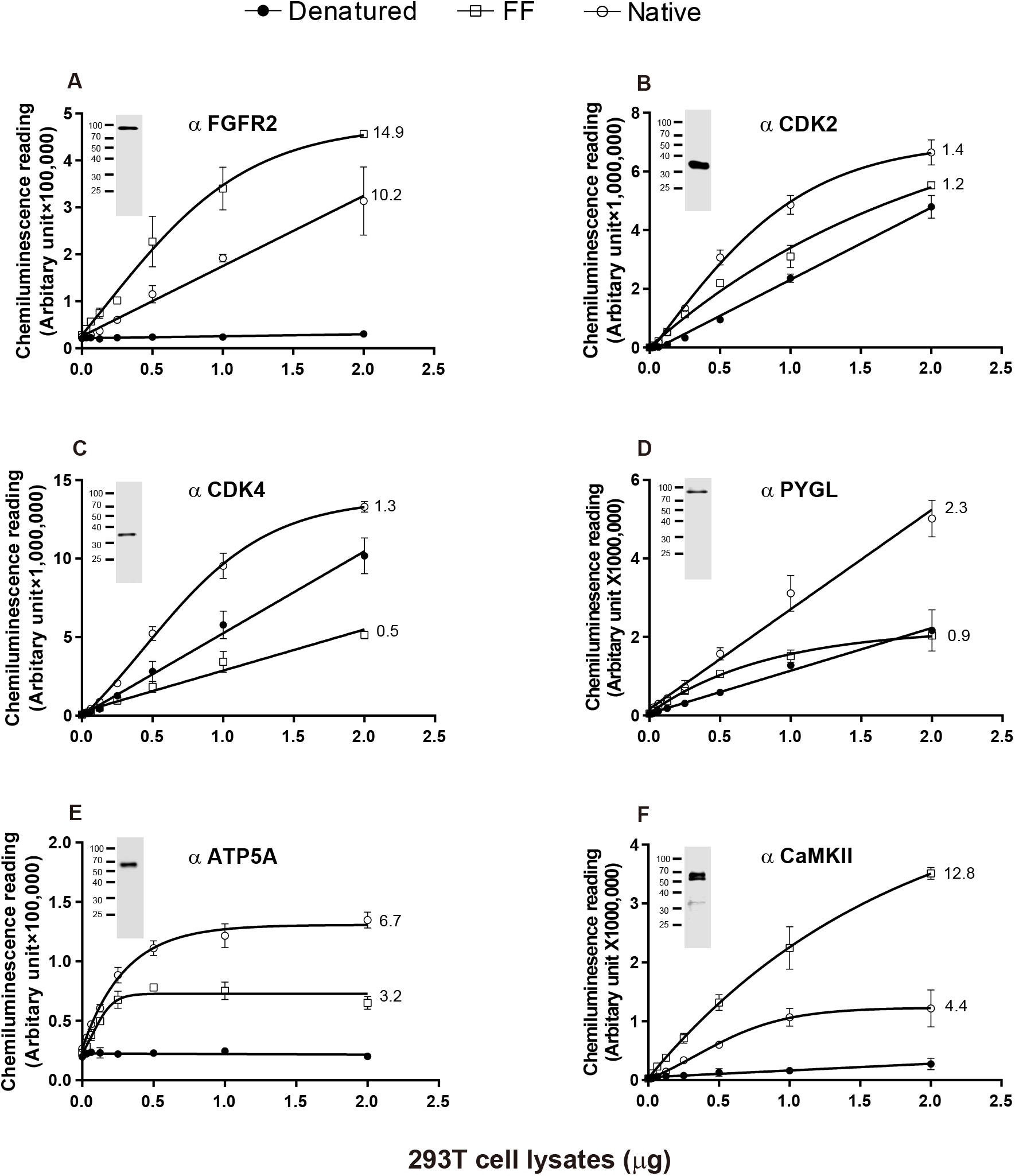
Comparison of performance of antibodies interacting with antigens at native, FF and denatured states in HEK293 cells. Whole cell lysates from HEK293 cells were extracted using lysis containing Triton-X 100 buffer, and used directly as native state, or treated with 1% SDS, and 50 mM DTT for 5 mins at 85°C as denatured state. In a parallel experiment, HEK293 cells were fixed in 10% Formalin Solution for 1 hour and cell lysates prepared as FF state. The total AAIs at native, denatured and FF states by each antibody was plotted as in 2A to 2F. The results from Western blot analysis of each antibody using denatured mouse liver lysates were inserted in 2A to 2F alongside.

The significantly increased AAIs at native state suggest that the contribution from conformation or shape dominates over that of the sequence of the epitope to the overall AAI at native state. Likewise, for many antigens at FF states, the contribution from shape is also the dominant factor to overall AAI. This observation naturally raises our concerns regarding the suitability of WB for validation of antibodies intended for ELISA and IHC analysis. In other word, the non-specific interaction observed with WB at denatured state may become negligible at native or FF states due to overwhelming contribution from the shape of the epitope to the overall AAI.

To test this hypothesis, we chose two polyclonal antibodies against different epitopes of PYGL. As shown in Fig. 3a, while WB revealed a strong band of expected size in mouse liver lysates with both antibodies, a band of smaller size was observed unexpectedly in mouse brain lysate when antibody-1, not antibody-2, was used. Clearly, antibody-1 would have been rejected as nonspecific for detecting PYGL in mouse brain lysate at denatured state.

**Fig. 3:**
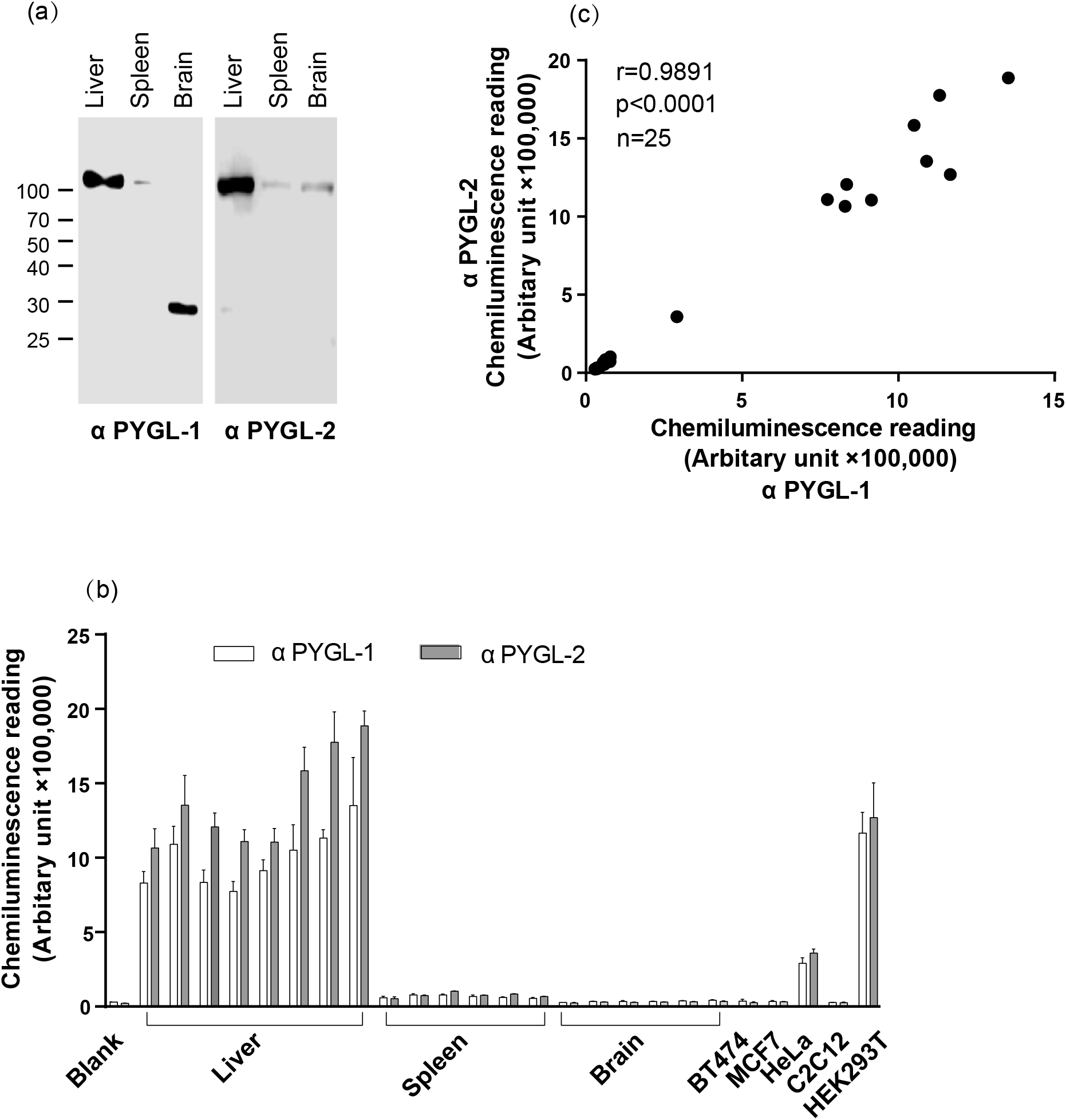
Cross-validation of the specificity of two anti-PYGL antibodies using protein lysates prepared from mouse liver, spleen, brain, and several cell lines at native state. a) Western blot analysis of PYGL protein levels in mouse liver, spleen and brain using two anti-PYGL polyclonal antibodies against different epitope of PYGL (PYGL-1 and PYGL-2). b) AAIs of PYGL-1 and PYGL-2 using total tissue lysates prepared from mouse liver, spleen and brain and several cell lines at native state. c) Correlation analysis of AAIs of PYGL-1 and PYGL-2 as described in Fig. 3b using Pearson’s correlation analysis.

Next, we measured overall AAIs of both antibodies using tissue lysates prepared from mouse liver, brain, spleen and several cellular lysates at native state using QDB method as shown in Fig. 3b. We found signals detected using these two antibodies are highly correlated with each other, with r=0.99 when analyzed with Pearson’s correlation analysis (Fig. 3c). Thus, our results suggested that both antibodies were highly specific for PYGL for mouse brain, liver, spleen and several cell lines at native state based on the proposed independent antibody strategy (2).

We interpreted our results as that the sequence is the sole discriminator of antibody specificity in WB. However, the shape of an antigen may become an additional and dominant discriminator at native and FF states to override any nonspecific interaction observed at denatured state. As a corollary, a detection antibody is more likely to be non-specific when analyzing samples at denatured state, when shape is no longer the key discriminator anymore. Consequently, WB may inadvertently eliminate many otherwise highly specific antibodies intended for ELISA and IHC in the past decades.

In fact, while WB is universally considered as a must for validation of antibodies intended for ELISA and IHC analyses (8, 9), some researchers indeed recognized that some antibodies performing well in WB may not necessarily perform well in IHC (9, 10). Conversely, some antibodies likely suitable for IHC analysis were also found to be nonspecific based on WB(11). Thus, the concept of context-dependent validation has been proposed to emphasize these phenomena in several commentaries (2, 12). However, this concept is primary based on empirically observations only. There is neither sound theoretical basis underlying this concept, nor does it provide any practical method to evaluate the magnitude of its influence on the specificity of an antibody. Our study, on the other hand, provided the first direct biochemical evidence to support this practice.

By examining the specificity of two anti-PYGL antibodies at native state, we also introduced QDB method as a convenient tool to validate the specificity of an antibody for IHC and ELISA analyses. Indeed, it is rather technically challenging to assess and validate the specificity of an antibody intended for IHC, considering the IHC result is expressed as an image. In our case, to perform the IHC analysis and quantify the resulted images of 25 specimens would be quite a daunting task for many researchers. However, as a “Do-it-Yourself” ELISA assay for lysates prepared at native, denatured or FF states, QDB method is demonstrated to be a convenient tool for this purpose in both basic research and clinical laboratories(5).

Additionally, in this study, all six antibodies were found to react more strongly to native antigen than denatured antigen, although the epitopes of their antigens would all be considered as “linear epitopes”. Clearly, the shape rather than the sequence of the epitopes is the dominant factor in AAIs even for “linear epitopes”. Thus, our results also raise the question regarding the suitability of the term “linear epitope” in literature for failing to recognize the contribution from shape to AAI even for continuous epitopes at native state. Indeed, it is hard to imagine the existence of an epitope without any conformational structure at native state. Accordingly, we propose to use “continuous epitope” instead of “linear epitope” to avoid the confusion in the future.

Caution should be warranted that while we were confident about the specificity of two PYGL antibodies, we were unsure about that of the rest of antibodies used in the study. While all other antibodies were confirmed as specific by WB, the drastically increased AAIs at native or FF states may also partially be derived from off-target binding. Nonetheless, this possibility is not going to affect our conclusions. Rather, it enforces our conclusion that the shape of the epitope contribute significantly to overall AAI. Our results thus questioned WB analysis as a suitable method to validate antibody intended for IHC or ELISA analyses.

In summary, using QDB method, we demonstrated that the conformation structure of the epitope contributes significantly to the overall AAI. In many cases, it can be the dominant determinant to overall AAI to alter the specificity of the antibody defined at denatured state. Consequently, WB analysis may unintentionally reject a large number of good antibodies specific for IHC and ELISA analysis for failing to recognize the contribution from the conformation structure of an epitope at native state. Thus, we propose to stop using WB analysis for validation of antibodies intended for IHC and ELISA analysis in future practice.

## Materials and Methods

### General reagents

All general reagents for cell culture were purchased from Thermo Fisher Scientific (Waltham, MA, USA). Protease inhibitors were purchased from Sigma Aldrich (St. Louis, MO, USA). Pierce BCA protein assay kit was purchased from Thermo Fisher Scientific (Calsband, CA, USA). All other chemicals were purchased from Sinopharm Chemicals (Beijing, P. R. China). QDB plate was provided by Quanticision Diagnostics, Inc (RTP, USA). Both HRP-labeled Donkey Anti-Rabbit and Anti-mouse secondary antibodies were purchased from Jackson Immunoresearch Lab (Pike West Grove, PA, USA). The primary antibodies used were anti-FGFR2 (rabbit monoclonal, ab109372, abcam, RRID unknown), anti-CDK2(rabbit polyclonal, sc-163, Santa Cruz, RRID: AB_631215), anti-CDK4(rabbit polyclonal, sc-260, Santa Cruz, RRID: AB_631219), anti-CaMKII (rabbit monoclonal, ab181052, abcam, RRID: AB_2891241), anti-ATP5A (rabbit monoclonal, ab176569, abcam, RRID:AB_2801536), anti-PYGL-1 (rabbit polyclonal, 15851-1-AP, Proteintech, RRID: AB_2175014, anti-PYGL-2 (rabbit polyclonal, DF12134, Affinity Biosciences, RRID: AB_2844939).

### Mouse tissues and Cell lines

The collection of mouse livers, mouse spleens and mouse brains was performed in accordance with the ethical review board of Binzhou Medical University (ER #2016-19) as described previously(3). The HEK293T, HeLa, C2C12, MCF7, BT474 cell lines were purchased from the Cell Bank of Chinese Academy of Sciences, Shanghai, China. All the five cell lines were cultured in DMEM culture media supplemented with 10% FCS and L-glutamine and harvested for cell lysates preparation.

### Formalin Fixation of mouse liver and HEK293 cells

Both HEK293T cell pellets and ∼ 50mg mouse liver tissues were incubated with 10% neutral buffered formalin for 1 hour and 24 hours respectively. Both tissue and cell pellets were rinsed 3 times with deionized water to remove residual formalin before lysis.

### Preparation of tissue and Cell line lysates

Protein lysates were prepared by lysing tissue and cell pellets in lysis buffer (50 mM Hepes, pH 7.4, 137 mM NaCl, 5 mM EDTA, 5 mM EGTA, 1 mM MgCl2, 10 mM Na2P2O7, 1% Triton X-100, 10% glycerol) supplemented with protease and phosphatase inhibitors (100 mM NaF, 0.1 mM phenylmethylsulfonyl fluoride, 5 μg/mL pepstatin, 10 μg/mL leupeptin, 5 μg/mL aprotinin) with a handhold tissue homogenizer for 1 minute before centrifugation at 12000 × g for 5 mins. The supernatant from each 1.5ml Eppendorf tube was collected for measurement of protein content using Pierce BCA protein assay kit. Protein lysates from frozen tissue and fresh cell pellets were defined as at native state. Those from Formalin Fixed tissue or cell pellets were defined as at FF state. A portion of native lysates were further treated by adding equal volume 2x Laemmli SDS sample buffer and heated at 85°C for 5 mins and it was defined as at denatured state.

### Western blotting

The denatured lysates of 20 μg/lane were loaded on 12% SDS–PAGE and transferred onto nitrocellulose filter membrane, then membrane was blocked with 4% nonfat milk in TBST(0.1% tween20, 0.1 M Tris–HCl, 0.5 M NaCl) for 1 hour at room temperature. The membrane was further incubated with primary antibody (anti-FGFR2: 1:1000, anti-CaMKII: 1:1000, anti-ATP5A: 1:500, anti-CDK2:1:1000, anti-CDK4: 1:1000, anti-PYGL1: 1:5000, anti-PYGL2:1:1000) for 2 hour, followed by washing membrane twice, each time for 5 mins used TBST before the membrane was incubated with the secondary antibody at 1:2500 for 1 hour. The membrane was washed with TBST three times, each time for 5 mins before ECL solution was added for signal detection using C-Digit (LI-COR Biosciences).

### QDB analysis

The QDB process was described in detail elsewhere with slightly modifications (3–5). In brief, the protein lysates of native, denatured and formalin fixed states were diluted serially as specified in the figure legends. The lysates were loaded at 2ul/unit in triplicate for QDB analysis. The plate was dried for 4 hour at room temperature, then blocked with blocking buffer (4% nonfat milk in TBST) for 1 hour. The plate was incubated with 100 μl/well primary antibody diluted in blocking buffer overnight at 4°C (anti-FGFR2: 1:1000, CaMKII: 1:1000, ATP5A: 1:500, CDK2: 1:1000, CDK4: 1:1000, PYGL-1: 1:5000, PYGL-2: 1:1000) and washed. The plate was incubated with anti-rabbit secondary antibody for 4 hours at room temperature before it was inserted into a white 96-well plate pre-filled with 100μl/well ECL reagent prepared according to the manufacture’s manual. The chemiluminescence signals were quantified using Tecan Infinite 200pro Microplate reader with the option “plate with cover”.

### Statistical analysis

All the statistical analyses were performed with GraphPad Prism7.0 software (GraphPad Software Inc., USA). The correlation used Pearson’s correlation coefficient analysis, p<0.05 was defined as statistically significant.

## Author contribution

YZ & WZ performed all the assays and data analysis, JZ designed and supervised the overall study and drafted the manuscript, MY contributed to data interpretation and edited the manuscript.

